# Boosting the biogenesis and secretion of mesenchymal stem cell-derived exosomes

**DOI:** 10.1101/2020.02.08.940122

**Authors:** Jinli Wang, Emily Bonacquisti, J. Nguyen

**Author notes:** Corresponding author Juliane Nguyen, PhD, Associate Professor, Division of Pharmacoengineering and Molecular Pharmaceutics, Eshelman School of Pharmacy, University of North Carolina at Chapel Hill.

## Abstract

A limitation to using exosomes to their fullest potential is their limited secretion from cells, a major bottleneck to efficient exosome production and application. This is especially true for mesenchymal stem cell (MSCs), which can self-renew but have limited expansion capacity, undergoing senescence after only a few passages, with exosomes derived from senescent stem cells showing impaired regenerative capacity compared to young cells. Here we examined the effects of small molecule modulators capable of enhancing exosome secretion from MSCs. Treatment of MSCs with a combination of methyldopamine and norepinephrine robustly increased exosome production by three-fold without altering the ability of the MSC exosomes to induce angiogenesis, polarize macrophages to the anti-inflammatory phenotype, or inhibit fibrosis. These small molecule modulators provide a promising means to increase exosome production by MSCs.

## Introduction

Exosomes have emerged as promising carriers for cancer vaccines and drugs, as vehicles and mediators of cardiac regeneration, and as possible alternatives to cell-based therapies [1]. For example, mesenchymal stem cell (MSC)-derived exosomes have been shown to mediate tissue regeneration in a variety of diseases including ischemic heart disease, lung injury, liver fibrosis, and cerebrovascular disease [2]. This is mainly due to their intrinsic regenerative capacity, in that they are able to induce angiogenesis, promote proliferation, prevent apoptosis, and inhibit inflammatory reactions [2]. In addition to their intrinsic biological functions, exosomes are also promising drug carriers due to their small size, excellent biocompatibility, and capacity to load specific and diverse therapeutic molecules including proteins, nucleic acids, and small molecules.

One limitation to using exosomes to their fullest potential is their limited secretion from cells, which constitutes a major bottleneck to efficient, large-scale exosome production. This is especially true for MSCs which, while able to self-renew, have limited expansion capability [3]. MSCs undergo senescence after only a few passages, and exosomes derived from senescent stem cells have impaired regenerative capacity compared to young cells [4]. MSCs have been transfected with lentiviruses carrying *MYC* to generate immortalized MSCs [3], but *MYC* is a proto-oncogene and thus could cause undesirable side effects if used therapeutically.

Therefore, enhancing exosome production is critical for both allogeneic and autologous exosome-based therapies because: (i) time is a limiting factor, since exosome-based therapies need to be administered as quickly as possible, particularly for cancer vaccination or to reduce the immunogenic effects caused by non-matching allogeneic cells; (ii) large quantities of exosomes are required for exosome-based therapies in a very short timeframe, and while upscaling of cell cultures is an option, this approach is limited by the growth kinetics and number of cells isolated from patients; and (iii) expanded exosome synthesis would allow the production of off-the-shelf therapeutics.

A recent screening approach in prostate cancer cells identified 23 significant exosome stimulators and inhibitors [5], with fenoterol, norepinephrine, forskolin, methyldopamine, and mephenesin the top five stimulators. However, the screen was limited to a single prostate cancer cell type, and the identified molecules have yet to be tested or shown to work in other cell types [5,6]. Given that manumycin-A, a microbial metabolite, inhibits exosome secretion in C4-2B castration-resistant prostate cancer cells but not in normal RWPE-1 cells, it is possible that small molecule modulators have different effects on different cell types [6].

Thus, the objectives of this study were to: (i) assess the effects of a set of small molecules shown to increase exosome production in prostate cancer cells on exosome production from MSCs; and (ii) determine if the small molecule modulators affected exosomal composition and the intrinsic regenerative capacity of MSC exosomes.

## MATERIALS AND METHODS

### Materials

Fenoterol hydrobromide and L-(–)-Norepinephrine-(+)-bitartrate were purchased from Sigma-Aldrich (catalog No. F1016 and 489350). Forskolin was purchased from Cayman Chemical (catalog No. 11018). N-Methyldopamine hydrochloride was purchased from Alfa Aesar (catalog No. J60306). Mephenesin was purchased from Ambeed (catalog No. A307218).

### Extracellular vesicle isolation

Exosomes were isolated from passage P1-P6 human bone marrow-derived MSCs obtained from the American Type Culture Collection (ATCC PCS-500-012) and cultured in MesenPRO RS™ Medium (GibcoTM, Gaithersburg, MD). After two days of MSC culture in EV-free medium, exosomes were isolated using differential centrifugation. Briefly, the cell culture medium was centrifuged at 2000 rpm for 10 min to remove cells and debris. Then, supernatants were centrifuged at 10,000 x g for 30 min at 4°C and the supernatants were transferred to ultracentrifuge tubes (Beckman Coulter, Brea, CA) and diluted with phosphate-buffered saline (PBS). Tubes were centrifuged twice at 100,000 x g for 70 min at 4°C with a Beckman ultracentrifuge (32Ti rotor from Beckman Coulter). Supernatants were removed and the pellets resuspended in phosphate-buffered saline (PBS) between centrifugations. Finally, the pellets were suspended in PBS for downstream analysis.

### Exosome characterization by nanoparticle tracking analysis (NTA)

The size distribution and concentration of exosomes were characterized by nanoparticle tracking analysis (NTA) as previously described [7,8]. Briefly, exosomes were diluted in PBS and measured three times in three different areas for each sample, with the average used to determine exosome concentration.

### Western blotting

Exosome pellets were re-suspended and lysed with RIPA buffer, incubated at 4°C for an additional 15 min for complete lysis, and combined with 4× LDS buffer. Samples were heated to 95°C for 5 min and then analyzed on a 4-12% gel (Bio-Rad Laboratories, Hercules, CA) using SDS running buffer. Transfer onto PVDF membrane was performed at 100 V for 120 min. Blots were incubated with anti-human CD63^+^ primary antibody (1:1000, Cat# 556019; Becton Dickinson, Franklin Lakes, NJ) or CD9 primary antibody (1:1000, Cat# 555370; Beckton Dickinson) overnight at 4°C and visualized using the Bio-Rad ChemiDoc MP Imaging System (Bio-Rad Laboratories).

### MTT assay

Cellular viability was analyzed using the MTT assay as previously described [7,9,10]. MSCs were seeded into 96-well plates and cultured overnight. Cells were treated with different concentrations of compounds for 24 h. MTT solution was added to the cells and incubated at 37°C for 3.5 h. Acidified isopropanol was added to dissolve the purple crystals, the plate was shaken for 15 min at RT, and the absorbance was read at 590 nm using an Epoch microplate spectrophotometer (BioTek, Winooski, VT). Background was removed by subtracting the absorbance recorded from a well containing no cells. Viability was calculated by dividing the absorbance from non-treated wells, which was set to 1.

### Fibrosis assay

Primary human cardiac fibroblasts were obtained from ScienCell (Cat# 6330, ScienCell Research Laboratories Inc., Carlsbad, CA). Passage number P3-P5 cells were taken, plated into 24-well plates, cultured in fibroblast medium 2 (Cat# 2331, ScienCell) containing 5% fetal bovine serum (FBS), and grown to 80% confluency. Cells were treated with 8 μg exosomes. After 12 h, cells were stimulated with an additional 500 μl of complete medium containing TGF-β (Peprotech, Rocky Hill, NJ) at a final concentration of 10 ng/ml. To assess collagen production, cells were lysed for RNA extraction after 48 h according to the TRIzol protocol (Thermo Fisher Scientific, Waltham, MA). cDNA was synthesized with the First Strand cDNA Synthesis Kit (New England BioLabs, Ipswich, MA) and analyzed for expression of collagen I (*COL1A1*) (primer sequences: forward GGGCAAGACAGTGATTGAATA and reverse ACGTCGAAGCCGAATTCCT), and *GAPDH* (forward CAAGGTCATCCATGACAACTTTG and reverse GTCCACCACCCTGTTGCTGTAG) by RT-PCR using the SYBR Supermix (Bio-Rad Laboratories) on a CFX Connect Real-Time PCR Detection System (Bio-Rad Laboratories).

### Angiogenesis assay

Primary human umbilical vein endothelial cells (HUVECs) were obtained from Gibco (Cat#: C0035C) and cultured in Medium 200 (Cat#: M200500, Thermo Fisher Scientific) containing large vessel endothelial supplement (Cat#: A1460801, Thermo Fisher Scientific). HUVECs were incubated with 7.5 μg of MSC-derived exosomes for 24 h. Then, HUVECs were plated into wells coated with Geltrex (Cat#: A1413201, Thermo Fisher Scientific) and cultured overnight. 17 h after plating, HUVECs were stained with calcein and imaged at em/ex 488/505 with a MiniMax Imager (Molecular Devices, Sunnyvale, CA). Angiogenesis was analyzed with ImageJ software using the angiogenesis analyzer.

### Bone marrow-derived macrophage (BMDM) isolation and culture

Bone marrow was harvested from the femurs and tibias of C57BL/6 mice as previously described [11]. After isolation and centrifugation, bone marrow cells were resuspended in DMEM/F12-10, frozen in DMEM/F12-40 with 10% DMSO and stored in liquid nitrogen until further use. Cells were thawed, washed once, and resuspended in macrophage complete medium (DMEM/F12, 10% FBS, 1% penicillin/streptomycin, 100 U/ml recombinant murine M-CSF; Peprotech, Rocky Kill, NJ; Cat# 315-02). 5 ml of macrophage complete medium was added on day 3. On day 7, cells were used for polarization assays.

### Macrophage polarization assay

BMDMs were treated with 16 μg/ml exosomes. 24 h after treatment, pro- and anti-inflammatory markers were quantified by RT-PCR. Briefly, cells were lysed with TRIzol reagent to extract RNA. cDNA was synthesized with the First Strand cDNA Synthesis Kit as above and analyzed for expression of *Inos* (forward CACCTTGGAGTTCACCCAGT, reverse ACCACTCGTACTTGGGATGC), *Il6* (forward ACTTCACAAGTCGGAGGCTT, reverse TGGTCTTGGTCCTTAGCCAC), *Arg1* (forward GTGAAGAACCCACGGTCTGT, reverse CTGGTTGTCAGGGGAGTGTT), *Cd206* (forward CAAGGAAGGTTGGCATTTGT, reverse CCAGGCATTGAAAGTGGAGT), and beta-actin (*Actb*) (forward GCCTTCCTTCTTGGGTATGG, reverse CAGCTCAGTAACAGTCCGCC) by RT-PCR using SYBR Supermix (Bio-Rad Laboratories) on a CFX Real-Time PCR Detection System (Bio-Rad Laboratories).

### Mechanism study

After 48 h compound treatment, MSCs were lysed and RNA were extracted with Trizol according to the manufacturer’s protocol (Invitrogen, Carlsbad, CA). cDNAs were synthesized using the ProtoScript® II First Strand cDNA Synthesis Kit (New England BioLabs) and analyzed for the expression of the exosome production-related genes listed in **Appendix Table 1** by RT-PCR on a CFX Connect Real-Time System (Bio-Rad Laboratories).

### Mass Spectrometric Analysis of Exosomes

MSCs were treated with norepinephrine and methyldopamine (NE + MeDA) for 24 h prior to switching to serum-free media for 24 h prior to exosome isolation. Exosomes were isolated using differential ultracentrifugation as described previously. EV samples were brought up to 30 μL in 1X LDS Sample Loading Buffer (Invitrogen), and then run into a 4-12% SDS-Page gel for a short period of time to remove contaminants. After staining with SimplyBlue SafeStain (Invitrogen), these regions were excised, cut into 1mm cubes, de-stained, then reduced and alkylated with dithiothreitol (DTT) and iodoacetamide (IAA), respectively (Sigma). Gel pieces were dehydrated with acetonitrile and digested with 10 ng/μL Trypsin in 50 mM ammonium bicarbonate for thirty minutes at room temperature, then placed at 37°C overnight. Exosomal peptides were extracted the next day by addition of 50% acetonitrile, 0.1% trifluoroacetic acid (TFA), then dried down in a CentriVap concentrator (Labconco). Peptides were desalted with homemade C18 spin columns, dried again, and reconstituted in 0.1% TFA.

Peptides were injected onto a 30 cm C18 column with 1.8 µm beads (Sepax), with an Easy nanoLC-1200 HPLC (Thermo Fisher), connected to an Orbitrap Fusion Lumos mass spectrometer (Thermo Fisher). Solvent A was 0.1% formic acid in water, while solvent B was 0.1% formic acid in 80% acetonitrile. Ions were introduced to the mass spectrometer using a Nanospray Flex source operating at 2 kV. Peptides were eluted off the column using a multi-step gradient that began at 3% B and held for 2 minutes, quickly ramped to 10% B over 6 minutes, increased to 38% B over 95 minutes, then ramped to 90% B in 5 minutes and was held there for an additional 3 minutes to wash the column. The gradient then returned to starting conditions in 2 minutes and the column was re-equilibrated for 7 minutes, for a total run time of 120 minutes. The flow rate was 300 nL/min throughout the run. The Fusion Lumos was operated in data-dependent mode with a cycle time of 2 seconds. The full scan was done over a range of 375-1400 m/z, with a resolution of 120,000 at m/z of 200, an AGC target of 4e5, and a maximum injection time of 50 ms. Peptides with a charge state between 2-5 were selected for fragmentation. Precursor ions were fragmented by collision-induced dissociation (CID) using a collision energy of 30 and an isolation width of 1.1 m/z. MS2 scans were collected in the ion trap with the scan rate set to rapid, a maximum injection time of 35 ms, and an AGC setting of 1e4. Dynamic exclusion was set to 45 seconds to allow the mass spectrometer to fragment lower abundant peptides.

### Bioinformatic Data Analysis

Raw data was searched with the SEQUEST search engine within the Proteome Discoverer software platform, version 2.2 (Thermo Fisher), using the UniProt human database. Trypsin was selected as the enzyme allowing up to 2 missed cleavages, with an MS1 mass tolerance of 10 ppm, and an MS2 mass tolerance of 0.6 Da. Carbamidomethyl on cysteine was selected as a fixed modification. Oxidation of methionine was set as a variable modification. Percolator was used as the FDR calculator, filtering out peptides with a q-value greater than 0.01. Label-free quantitation was performed using the Minora Feature Detector node, with a minimum trace length of 5. The Precursor Ions Quantifier node was then used to calculate protein abundance ratios, using only unique and razor peptides. The pairwise based method was employed to calculate the protein ratios, which uses a protein’s median peptide ratio to determine the protein ratio. Normalization was done using the Total Peptide Amount method. Comparison of the treated and non-treated groups was analyzed using Graphpad Prism 8. Protein pathway analyses were conducted using a combination of KEGG, Reactome, and Panther gene hits. Protein interactors were determined utilizing StringDB.org and uploaded into Cytoscape 3.2.1, before being fed into ClueGo 2.5.4 for visualization. Proteins that did not fit these parameters were removed from subsequent analyses due to lack of interaction evidence. Only gene hits with statistical significance of P<0.05 with a Bonferroni step-down correction were considered for further discussion.

## Results

### Effects of small molecules on exosomal production efficiency

Five compounds were selected to assess if they could stimulate MSCs to enhance their exosome production: four FDA-approved drugs (fenoterol, norepinephrine, methyldopamine, and mephenesin) and the other a nutritional supplement (forskolin). To determine a range of non-toxic concentrations, MSCs were treated with the five different compounds at concentrations between 10 and 100 µM. Two independent assays were used to determine the effects of the compounds on cells: (a) cell and metabolic activity as determined by the MTT assay, and (b) total cell number. Treatment of MSCs with these compounds at the tested range of concentrations did not affect MSC proliferation and had no obvious cytotoxicity (**Fig. 1A and 1B**).

**Figure 1.**
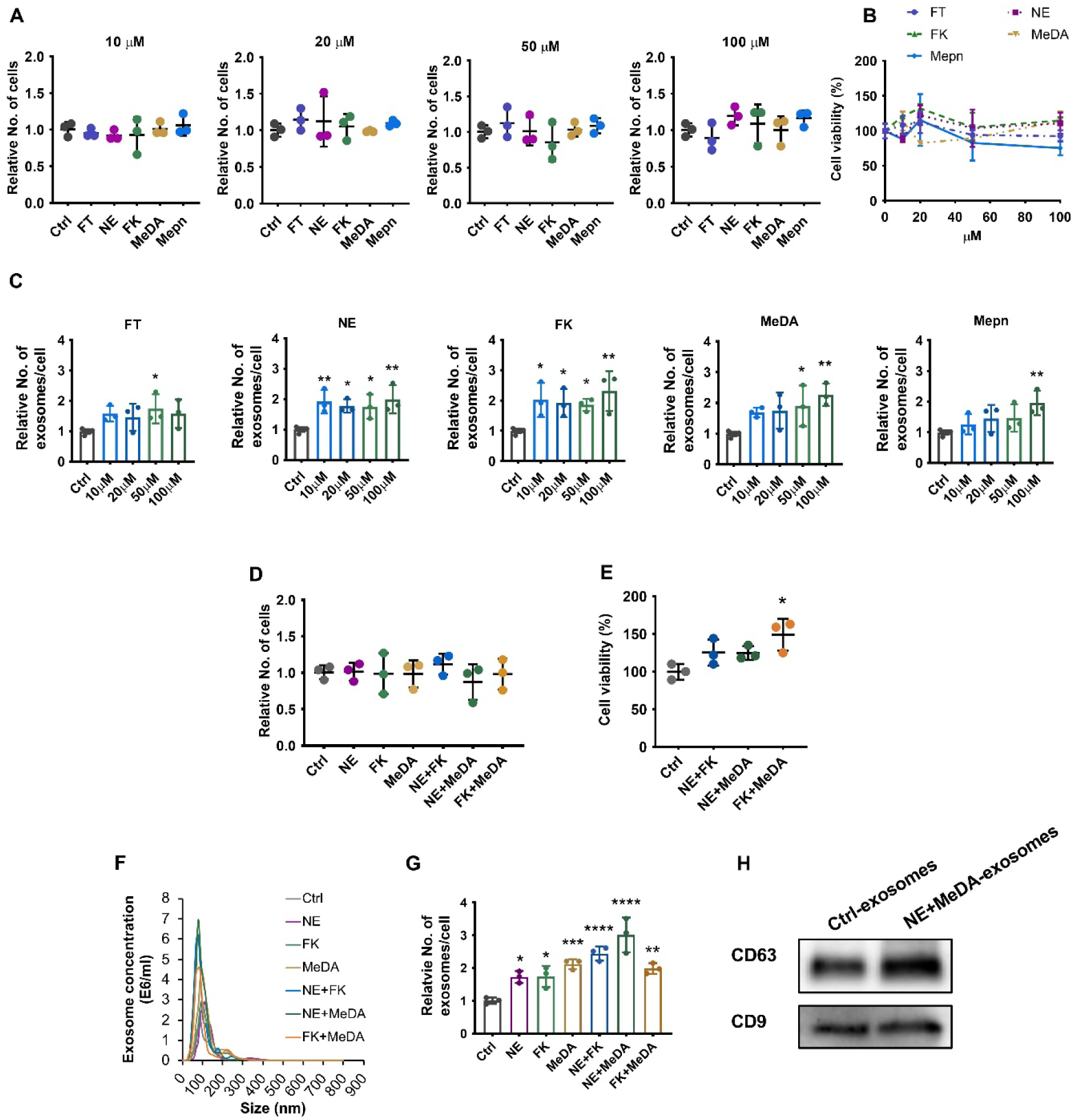
Effects of small molecules on MSC cell viability and exosome production efficiency. (**A**) Cell proliferation rate. (**B**) Cell viability and metabolism measured by the MTT assay. (**C**) Production efficiency of exosomes derived from MSCs treated with different doses of compounds. (**D**) Cell proliferation rate of MSCs in response to treatment with small molecule modulators. (**E**) Cell viability and metabolism of MSCs after treatment with small molecule modulators measured by the MTT assay. (**F**) Size distribution of exosomes by NTA in response to treatment of MSCs with small molecule modulators. (**G**) Production efficiency of exosomes derived from different MSCs treated with combinations of compounds. (**H**) CD63 and CD9 exosomal surface marker protein expression by western blotting. One-way ANOVA with Dunnett’s multiple comparisons test; * P<0.05, **P<0.01, ***P<0.001, ****P<0.0001, n=3-6 each group. FT: fenoterol; NE: norepinephrine; FK: forskolin; MeDA: methyldopamine; Mepn: mephenesin.

After confirming that all five compounds were well tolerated by MSCs even at high concentrations, the effects of these small molecules on MSC exosome production efficiency were examined. All the tested doses of norepinephrine (NE) and forskolin (FK) increased exosome production significantly and by similar levels (1.75-to 2-fold for norepinephrine and 1.9-to 2.3-fold for forskolin, P<0.05). 50 µM fenoterol (FT) increased exosome production by approximately 1.7-fold, P<0.05). Methyldopamine (MeDA) and mephenesin (Mepn) increased exosome production in a dose-dependent manner, with 100 µM showing the greatest effect on exosome production (∼2.3-fold for MeDA, P<0.001; and ∼2-fold for Mepn, P<0.01) (**Fig. 1C**).

Combining norepinephrine with forskolin and norepinephrine with methyldopamine further enhanced exosome production by ∼2.5-fold and ∼3-fold, respectively (**Fig. 1G**). Adding forskolin to methyldopamine did not further increase exosome production (**Fig. 1G**). While the combination treatment did not affect overall cell counts (**Fig. 1D**), they increased metabolic activity of MSCs by ∼1.5-fold as measured by the MTT assay (**Fig. 1E**).

Exosomes derived from non-treated MSCs and MSCs treated with the small molecule compounds measured 80 to 100 nm and not differ in size (**Fig. 1F**). The isolated exosomes expressed the exosomal surface marker proteins CD63 and CD9 (**Fig. 1H**), confirming their identity as exosomes.

### Anti-fibrotic effects

Exosomes derived from MSCs have been shown to have intrinsic anti-fibrotic effects. To determine if compound treatment affected the intrinsic biological effects of the produced MSC exosomes, cardiac fibroblasts were incubated with exosomes derived from untreated and treated MSCs.

To induce collagen expression, cardiac fibroblasts were stimulated with TGF-β (**Fig. 2A**). Untreated MSC-derived exosomes significantly decreased TGF-β-induced collagen expression from 2.3-fold to ∼1.5-fold (P<0.001). All the compound-treated MSC-derived exosomes decreased TGF-β-induced collagen expression, indicating that compound-treated MSC-derived exosomes preserve their intrinsic anti-fibrotic effects (**Fig. 2A**). The MTT assay showed that MSC-derived exosome did not affect fibroblast cell viability, indicating the anti-fibrotic effects of exosomes were not secondary to cytotoxicity (**Fig. 2B**). We further tested the direct effect of the compounds on fibrosis and found that MeDA and combined compound treatment significantly inhibited TGF-β-induced collagen expression but not cell viability (**Fig. 2C-D**). These results suggest that MeDA and compound combinations also inhibit fibrosis.

**Figure 2.**
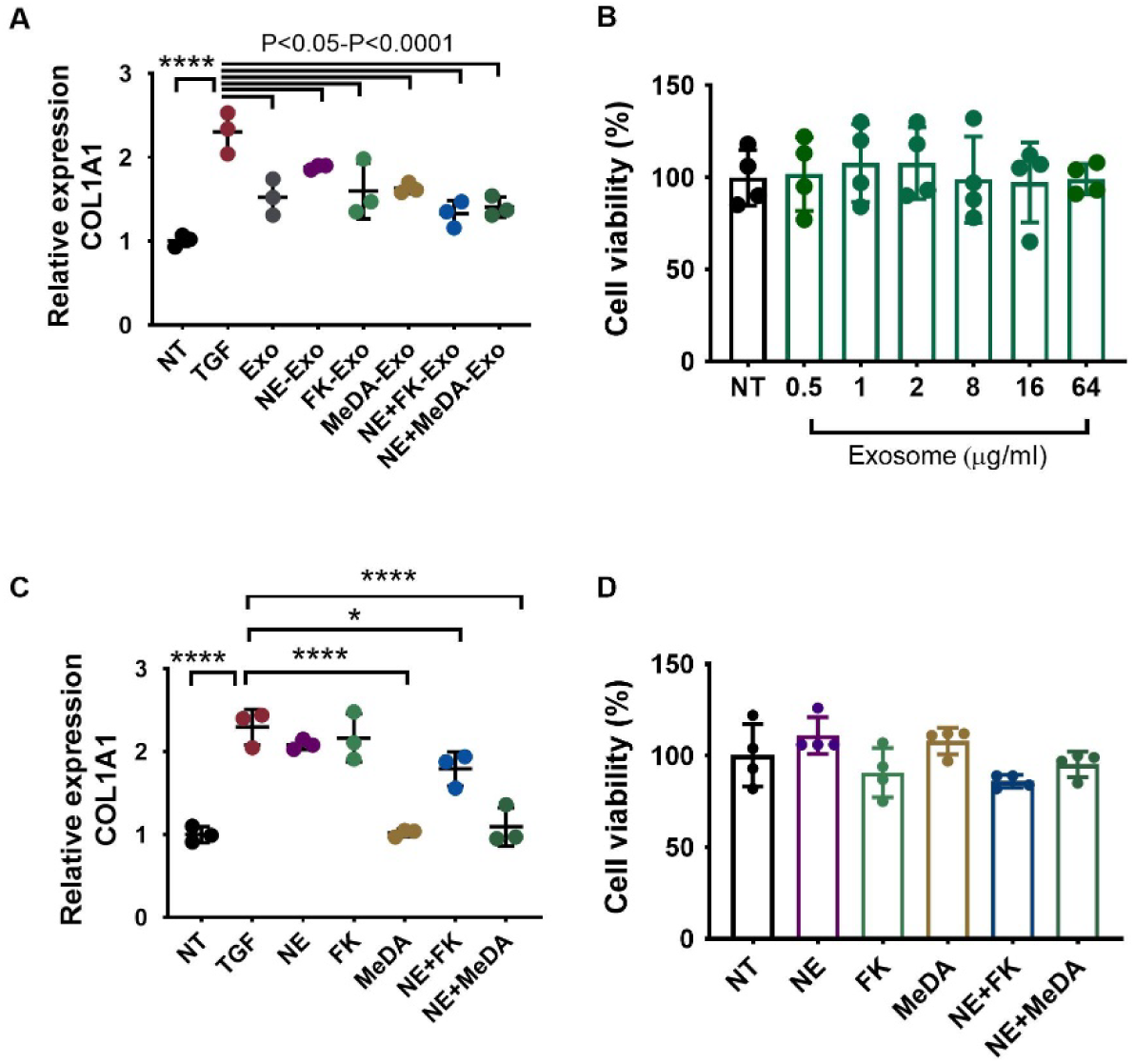
Anti-fibrotic effects of exosomes derived from MSCs treated with small molecule modulators. (**A**) *COL1A1* expression in human cardiac fibroblasts (HCF) incubated with exosomes isolated from MSCs treated with small molecule modulators as quantified by RT-PCR. (**B**) MTT assay of MSC-derived exosomes on HCFs. (**C**) *COL1A1* expression in human cardiac fibroblasts (HCF) incubated with small molecule modulators. (**D**) MTT assay of compound treatment on HCFs. One-way ANOVA with Dunnett’s multiple comparisons test, * P<0.05, **P<0.01, ***P<0.001, ****P<0.0001; n=3-4.

### Macrophage polarization

In addition to their anti-fibrotic effects, MSC-derived exosomes have anti-inflammatory effects and polarize macrophages to the anti-inflammatory M2 phenotype. As shown in **Fig. 3**, MSC-derived exosomes had negligible effects on the M1 marker *iNOS* but significantly decreased *IL6* expression compared to untreated cells (NT group). Similar to exosomes derived from untreated MSCs, exosomes derived from MSCs treated with the small molecules did not alter *iNOS* expression but significantly decreased *IL6* levels (**Fig. 3A**). Except for the forskolin-treated group, exosomes derived from non-treated MSCs and MSCs treated with norepinephrine and methyldopamine increased anti-inflammatory M2 markers *Arg1* and *CD206* (**Fig. 3B**). Almost all the MSC-derived exosomes increased the M2/M1 ratio, but those induced by forskolin failed to significantly increase *Arg1*/*iNOS, Arg1*/*IL6*, and *CD206*/*IL6* (**Fig. 3C**). These results suggest that treatment of MSCs with the compounds did not alter the intrinsic anti-inflammatory effects of exosomes.

**Figure 3.**
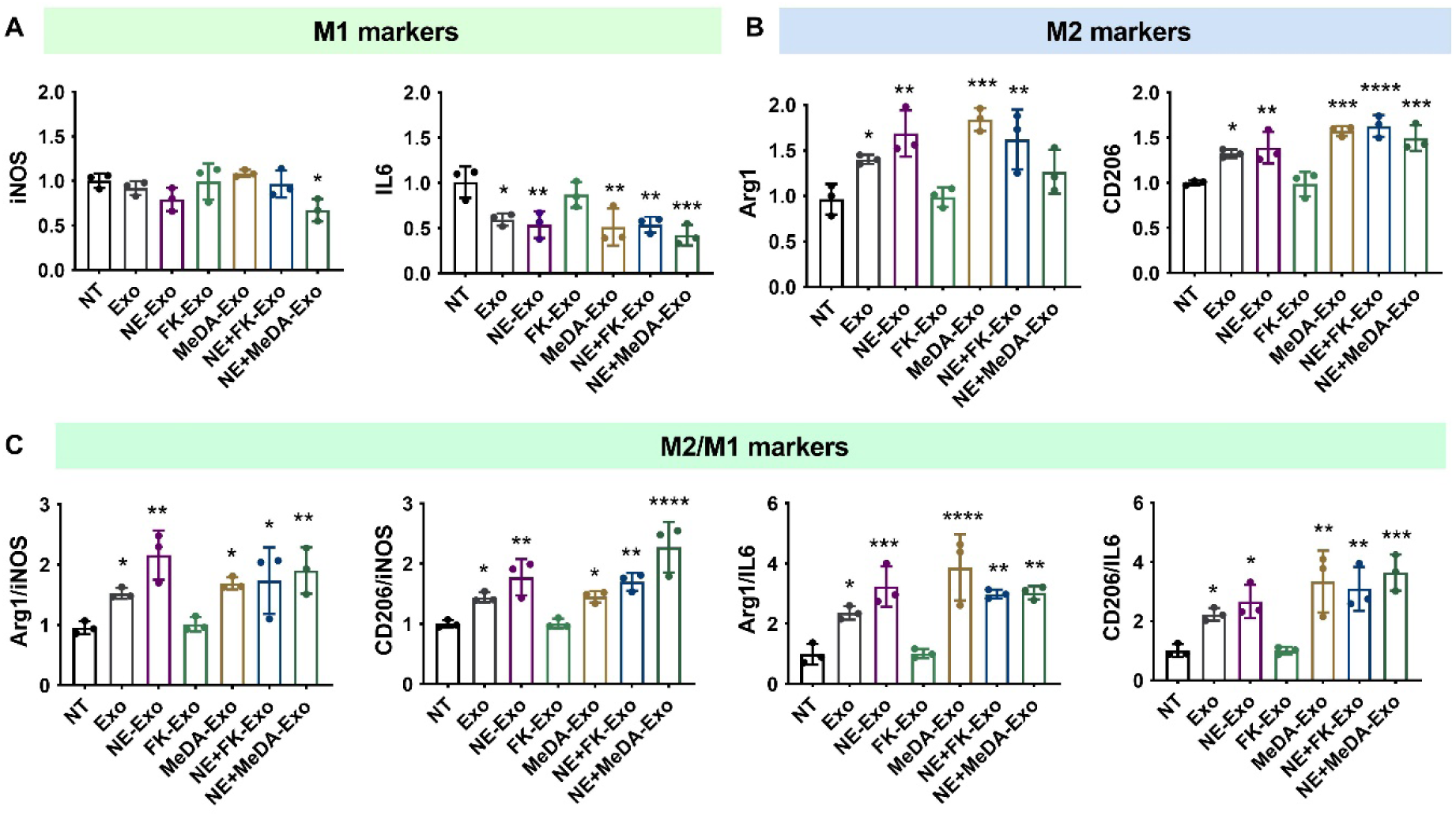
Polarization of macrophages with compound-treated MSC-derived exosomes. (**A**) Inflammatory macrophage (M1) markers *iNOS* and *IL6*. **(B)** Anti-inflammatory macrophage (M2) markers *Arg1* and *CD206*. (**C**) M2/M1 ratios. One-way ANOVA with Dunnett’s multiple comparisons test, * P<0.05, **P<0.01, ***P<0.001, ****P<0.0001; n=3.

### Angiogenesis effect

To assess if compound-induced exosomes had altered ability to induce angiogenesis, HUVECs were incubated with exosomes derived from untreated MSCs and MSCs treated with the small molecules. Similar to HUVECs treated with exosomes derived from untreated MSCs, exosomes derived from treated MSCs significantly increased parameters for tube formation such as increased total length, total branching length, number of junctions, number of nodes, number of meshes, and mesh index (**Fig. 4**). Exosomes induced by small molecules preserved their intrinsic pro-angiogenic function.

**Figure 4.**
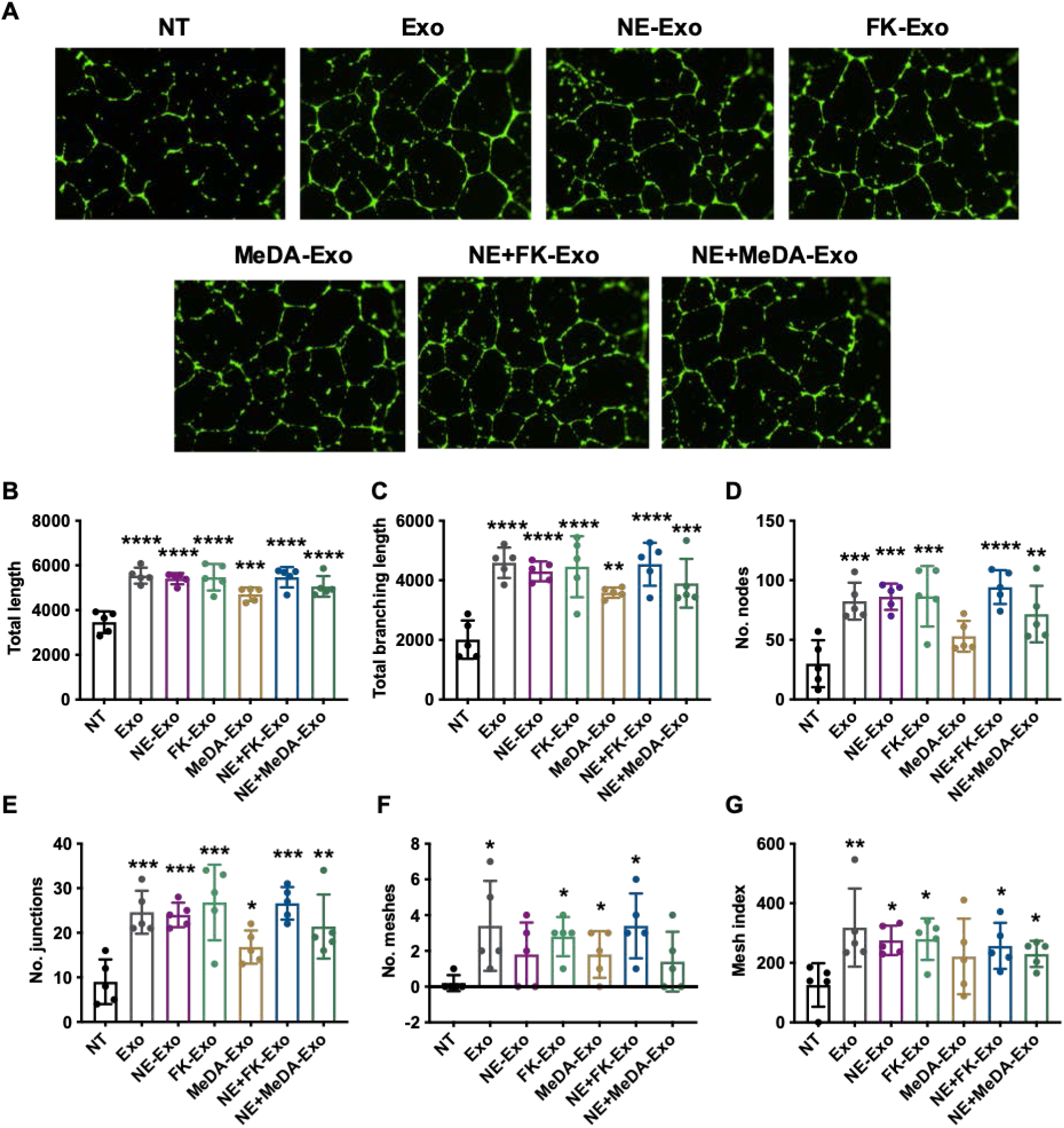
Tube formation assay. **(A)** Representative images of the angiogenesis effect of compound-treated MSC-derived exosomes. **(B-G)** Quantification of angiogenesis using ImageJ. **(B)** Total length**. (C)** Total branching length. **(D)** Number of nodes. **(E)** Number of junctions. **(F)** Number of meshes. **(G)** Mesh index. One-way ANOVA with Dunnett’s multiple comparisons test, * P<0.05, **P<0.01, ***P<0.001, ****P<0.0001; n=5.

### Effects of small molecule modulators on genes affecting exosome biogenesis and secretion

Cellular exosome production involves several processes that include the inward budding of the late endosomal membrane to form intraluminal vesicles within multivesicular bodies (MVBs). This is followed by fusion of MVBs with the plasma membrane prior to exosomal release into the extracellular milieu [12].

We first assessed the effects of the small molecule antagonists on neutral sphingomyelinase (nSMase2; *SMPD3*) expression, because it has been shown to be a key lipid metabolic enzyme in early exosome formation [12]. Ceramide is one of the main regulators of exosome formation and it is formed after the hydrolytic removal of the phosphocholine moiety of sphingomyelin by sphingomyelinase. Ceramide triggers the inward budding and formation of intraluminal vesicles resulting in multivesicular bodies [13]. Treatment of MSCs with small molecule modulators significantly upregulated nSMase2 by 1.5-to 2-fold (**Fig. 5A**). In contrast, the small molecule modulators did not affect expression of *Hrs, Tsg101, Stam1*, and *Alix*, which are known to regulate the ESCRT-dependent biogenesis of exosomes (**Fig. 5B-E**) [14].

**Figure 5.**
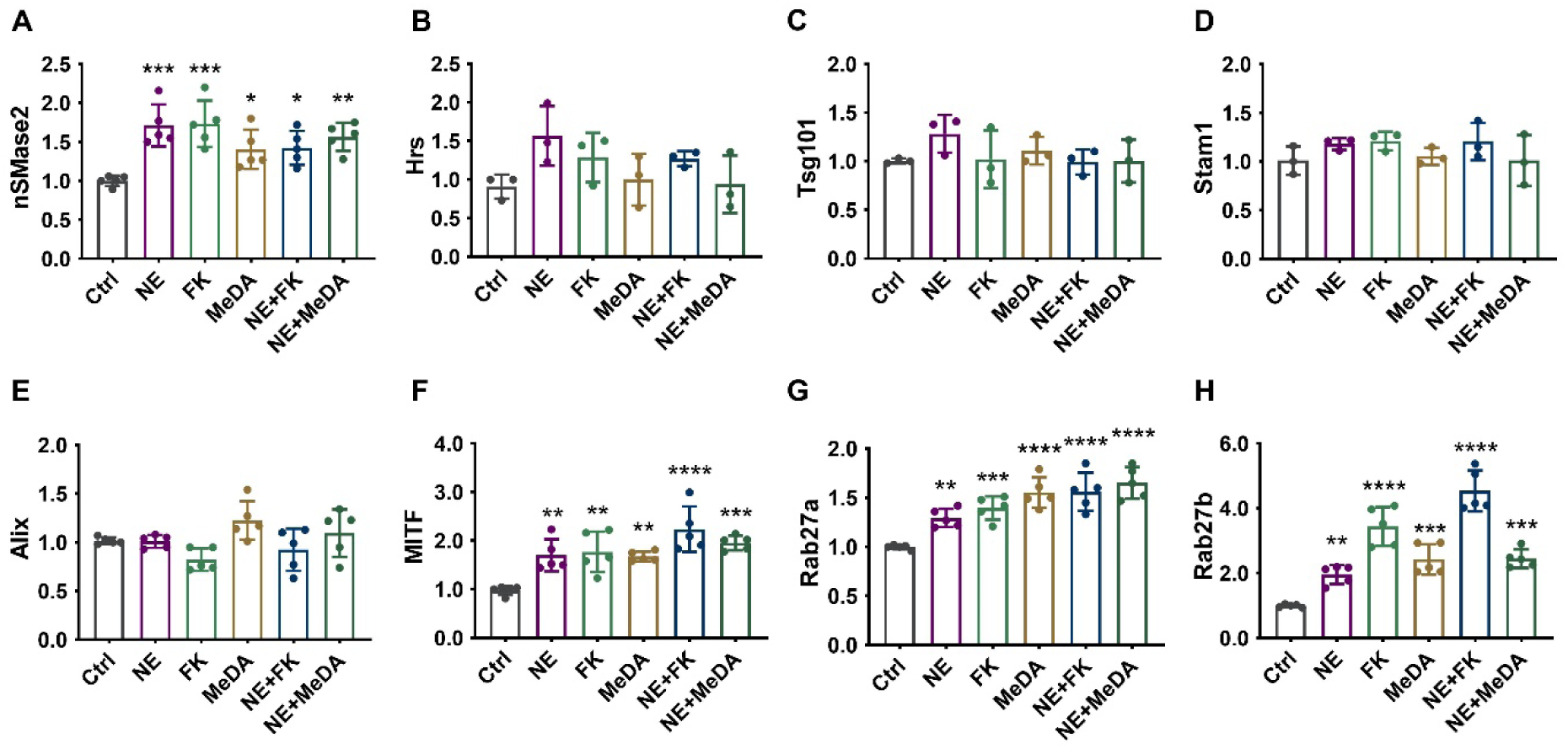
The effect of small molecule compounds on exosomal production-related pathways after 48 h treatment. RT-PCR quantification of **(A**) nSMase2 (*SMPD3*), **(B)** *Hrs*, **(C)** *Tsg101*, **(D)** *Stam1*, **(E)** *Alix*, **(F)** *MITF*, **(G)** *Rab27a*, and **(H)** *Rab27b*. One-way ANOVA with Dunnett’s multiple comparisons test, * P<0.05, **P<0.01, ***P<0.001, ****P<0.0001; n=3-5.

Next, we assessed the effects of the small molecule modulators on microphthalmia-associated transcription factor (*MITF*). *MITF* has been reported to play an important in exosome generation and is known to increase the expression of late endosomal proteins such as Rab7 and Rab27a, which are key proteins in exosomal biogenesis and secretion [15,16]. The small molecule modulators significantly upregulated *MITF* by 1.5-to 2-fold (**Fig. 5F**).

Finally, we assessed how the small molecule modulators affected *Rab27a* and *Rab27b*. Rab family proteins have been reported to promote exosomal transportation and secretion. Rab27a/b have also been reported to be required for (a) MVB distribution to the cell periphery, and (b) MVB docking at the plasma membrane for exosome exocytosis [17]. Upon stimulation with small molecule modulators, *Rab27a* and *Rab27b* were increased ∼1.3 to 5-fold (**Fig. 5G&H**).

### Effects of small molecule modulators on exosomal protein composition

Because small molecule treatment might not only modulate exosome secretion but could also affect exosome composition, proteomics analysis was performed on exosomes derived from untreated MSCs and MSCs treated with a combination of norepinephrine and methyldopamine, which had the greatest effect on exosome secretion. In order to detect differences in protein abundance, proteomic data were analyzed using global and individual statistical comparisons. There were no statistically significant differences in total protein composition and amounts between exosomes derived from unmodified MSCs and exosomes derived from MSCs treated with norepinephrine and methyldopamine (**Fig. 6A and 6B**). Of the 2159 proteins identified in each sample, 7 (0.3%) were statistically significantly downregulated. 14 (0.6%) of the total protein content was statistically significantly higher in abundance (**Fig. 6C, Appendix Fig. 2**). Each of these proteins was enriched between ∼2 and ∼5 fold over the untreated.

**Figure 6.**
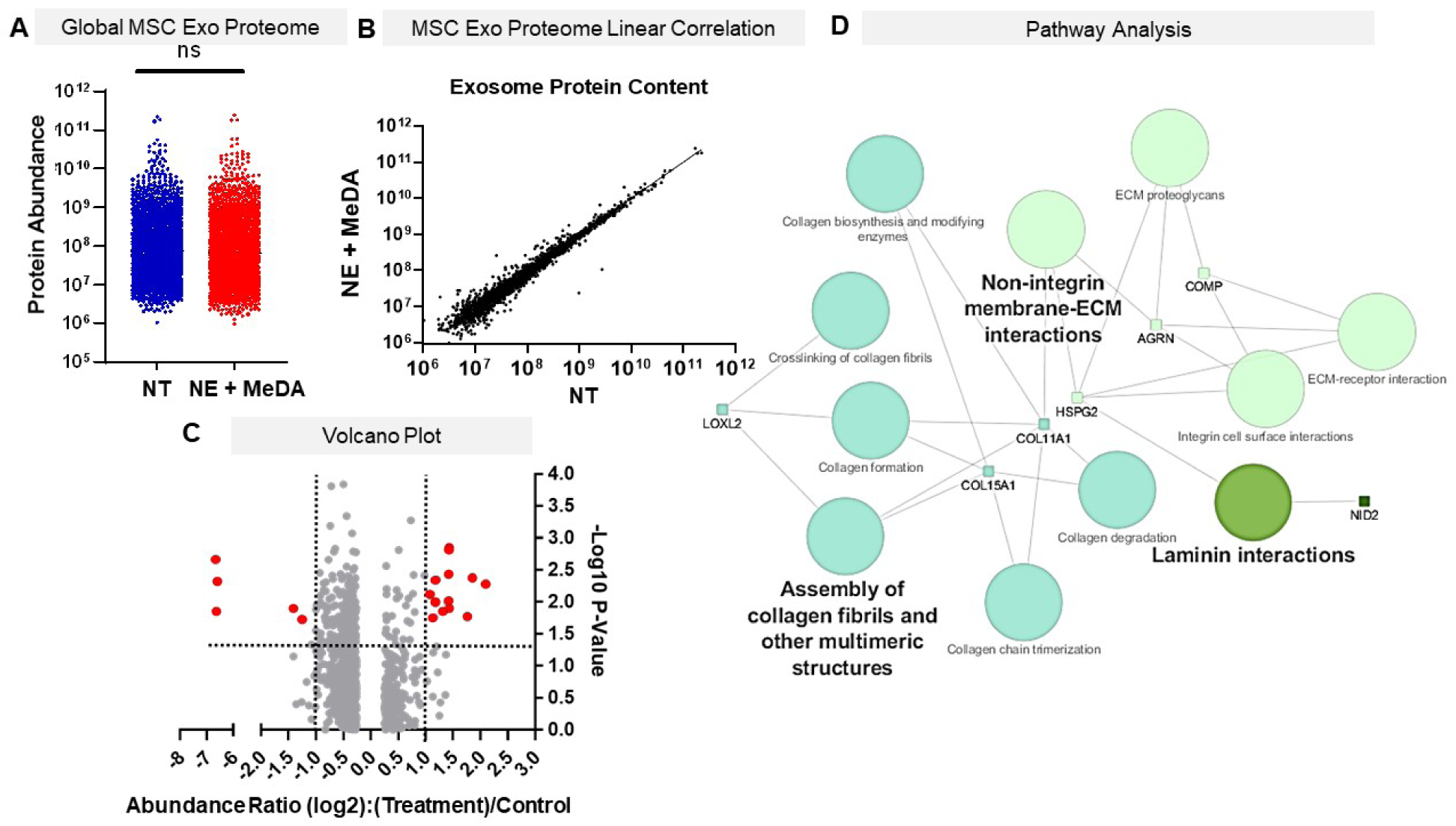
Proteomic profile of MSC exosomes 48h after NE + MeDA Treatment. (**A**) Raw abundance values from the NT and NE +MeDa groups showed no statistical difference using a paired t-test across samples. (**B)** Linear regression analysis comparing non-treated MSCs (NT) and MSCs treated with NE + MeDA. (**C**) Statistical analysis was performed by means of an unpaired two-tailed t-test of 14 statistically significant differentially expressed proteins with * P < 0.5, **P < 0.01, 2159 proteins per replicate. Data visualized using a volcano plot with red dots indicating statistically significantly different proteins with an abundance change of 2-fold or higher. (**D**) Cytoscape 3.2.1 visualization of Kegg, Reactome, and Panther Pathway hits of differentially expressed proteins using ClueGo v.2.5.4. Font size mapped to P value with Bonferroni step down correction, node size mapped to number of statistically significant gene hits.

control. Of these 14, only seven (LOXL2, COL15A1, COL11A1, HSPG2, AQRN, NID2 and COMP), were upregulated above the cumulative average protein abundance (**Appendix. Fig. 2**). None of the downregulated proteins were at or above the cumulative average, so this along with the lack of statistical significance excluded them from pathway analysis (**Appendix. Fig. 2**). The combinatorial pathway and interaction analysis was stringently annotated and analyzed with the pathway visualization software ClueGo. Three pathway databases, Panther, KEGG, and Reactome were used for analyses in order to create a comprehensive pathway visualization. Statistically significant pathways are visualized in **Figure 6D**, with the font and node size corresponding to statistical significance, and number of pathway gene hits, respectively. The pathways that were most represented were assembly of collagen fibrils and other multimeric structures, laminin interactions, and non-integrin membrane-ECM interactions.

## Discussion

Exosomes derived from MSCs are known to have intrinsic tissue regenerative effects; that is, they induce angiogenesis and cellular proliferation and inhibit fibrosis [2]. However, their widespread therapeutic use as drug carriers is currently hampered by their limited cellular secretion [18]. Exosome yield using routine cell culture is generally low and could significantly slow production, which is of particular concern if exosomes need to be derived rapidly from the patient’s own cells for personalized therapy. Having the means to significantly increase exosome secretion will be a highly important tool for the development of exosomes as drug carriers and their related clinical applications.

To date, upscaling of exosome production has involved increasing the volume of cell culture from flasks to containers, bioreactors, or hollow fibers [18,19]. These methods require substantial volumes and are impractical for large-scale production. Thus, alternative complementary or stand-alone methods to existing approaches would be highly useful.

GW4869 and manumycin-A were the first and few molecules known to inhibit exosome secretion [6,20]. Recently, prostate cancer cells were screened with small molecules to test their effects on exosome production [5,6], but the molecules identified in that screen (tipifarnib, nitrefazole, pentetrazol, etc.) have not been tested or shown to work in other cell types [5], despite evidence of cell type-specific effects of small molecules on exosome secretion [6]. Here we assessed the effects of a set of small molecules on exosome production from MSCs.

Five compounds (fenoterol, norepinephrine, forskolin, methyldopamine, and mephenesin) were selected from a screen previously reported to enhance exosome secretion in C4-2B castration-resistant prostate cancer cells [5]. While these compounds induced a 3.5-to 5.7-fold increase in exosome secretion in C4-2B prostate cancer cells at 10 µM concentrations, fenoterol, methyldopamine, and mephenesin failed to enhance exosome secretion at low doses in MSCs and only slightly enhanced exosome secretion at significantly higher doses (50 µM for fenoterol, 100 µM for mephenesin, and 50 µM for methyldopamine). Norepinephrine and forskolin were reported to increase exosome secretion in C4-2B cells by 4.6- and 5.7-fold. When tested in MSCs, exosome secretion was enhanced by only two-fold. When dual combinations of the small molecules were tested, norepinephrine co-dosed with methyldopamine had the greatest effects on exosome secretion in MSCs with a three-fold enhancement. This strongly suggests that exosome secretion is a cell-specific process and that small molecule modulators do not necessarily have the same effects on all cell types [5,6].

We showed that enhanced exosome secretion is not due to an increase in cell number after small molecule modulator treatment but could be the result of enhanced metabolic activity [21]. This is consistent with our target prediction analysis that revealed that forskolin mainly binds to proteins related to energy (ATP and glucose) production (**Appendix Fig.1 and Table 2**). Methyldopamine binds to the dopamine and adrenergic receptors, and norepinephrine binds to adrenergic receptors, whose activation has been shown to increase cellular metabolism (**Appendix Fig.1 and Table 2**) [22].

We chose three representative assays to assess if the small molecule modulators affected the intrinsic regenerative effects of MSC exosomes [8]. Exosomes derived from MSCs treated with small molecule modulators retained their antifibrotic effects and showed similar inhibition of collagen in cardiac fibroblasts as the exosomes derived from unmodified MSCs. Similarly, there were no differences in their ability to polarize macrophages to the M2 anti-inflammatory phenotype. Finally, in a tube formation assay, MSC exosomes derived from MSCs treated with small molecule modulators showed comparable angiogenic ability as control exosomes. This strongly indicates that chemically induced exosomes preserve their biological functions. In line with this, small molecule treatment of MSCs did not dramatically affect exosomal protein composition, in agreement with findings where an increase in Rab27a and Rab27b in HeLa cells did not affect exosomal protein composition [23]. The increased metabolic activity of the MSCs after small molecule treatment remained independent of exosome protein biogenesis, further showing the utility of the treatment as a powerful biochemical tool for increasing the throughput of exosome research.

With respect to specific pathways, the most potent small molecule modulators increased exosome secretion in MSCs by enhancing expression of genes related to the ESCRT-independent pathway, such as nSMase2 [24-26]. nSMase2 is a key regulator of MVB budding from early endosomes by generating ceramide. Ceramide is essential for exosome formation [13]. Additionally, the small molecule modulators upregulated other genes such as MITF, Rab27a, and Rab27b, which are known to be essential for the transportation and secretion of exosomes [23,27]. The pathways modulated by the eight upregulated proteins support observations from several groups, as exosomes have been heavily implicated in altering and binding to ECM proteins to promote cell migration and cell adhesion [28-30]. Assembly of collagen fibrils, as well as integrin dependent and independent binding are both necessary for propagation of wound healing responses after injury. Bioinformatic pathway analyses also yielded gene hit counts against laminin-dependent pathways, further supporting our *in vitro* data, as laminins play a critical role in reepithelization and angiogenesis [31]. The up-regulated proteins used for pathway prediction analysis are directly in line with the angiogenesis, macrophage, and tube formation assay, as they are all involved with exosomal binding to ECM proteins and promotion of wound-healing effects [32-34]. Despite these proteins only composing 0.6% of the total protein sample, they are highly enriched (significantly above average protein abundance) and could be of therapeutic benefit in tissue regeneration and wound healing.

## Conclusion

We have shown that exosome secretion in MSCs can be enhanced by treatment with small molecule modulators. These small molecule modulators did not affect the intrinsic regenerative effects of MSC exosomes and did not significantly alter total exosomal protein expression levels. The proteomic alterations that were observed appear to potentially enhance the MSC exosomes’ therapeutic action. Thus, these compounds could be used to enhance exosome secretion from MSCs for practical application. Further investigation into the structural optimization of small molecule modulators capable of enhancing exosome production is warranted.

## Author Contributions

Conceptualization: J.N.; Methodology: J.N. and J.W.; Software, Graphpad Prism; Formal Analysis, J.W., E.B. J.N.; Experiments and Data Curation, J.W & E.B.; Writing – Original Draft Preparation, J.W. and J.N.; Writing – Review & Editing, E.B., J.N; Supervision, J.N.; Project Administration, J.N.; Funding Acquisition, J.N,

## Acknowledgements

We thank Kevin Welle at Rochester University (URMC Mass Spectrometry Resource Facility) for performing the mass spectrometry data acquisition. We thank Dr. John Canty for generously allowing us to use the facilities and equipment at CTRC.

## Abbreviations

MSC: mesenchymal stem cell
FT: Fenoterol
NE: norepinephrine
FK: foskolin
MeDA: methyldopamine
Mepn: Mephenesin
HCF: human cardiac fibroblast
HUVEC: human umbilical vein endothelial cell
BMDM: bone-marrow derived macrophage
MVB: multi-vesicular body
ESCRT: endosomal sorting complex required for transport

