## Supplementary File for "Boosting the biogenesis and secretion of mesenchymal stem cell-derived exosomes"

**Appendix**

**Appendix Table 1: Primer sequences for RT-PCR**

| No. | Gene | 5'-3' forward primer | 5'-3' reverse primer |
| --- | --- | --- | --- |
| 1 | nSMase2 | CAACAAGTGTAACGACGATGCC | CGATTCTTTGGTCCTGAGGTGT |
| 2 | Hrs | CTCCTGTTGGAGACAGATTG | CAGGTACAGGATCTTGTTAC |
| 3 | Tsg101 | GAGAGCCAGCTCAAGAAAATGG | GGGATTGTTCCAGTGAGGTTC |
| 4 | Stam1 | GATGAATACTGCTGAGGACT | CTGAGAGCCAATAGCTGGGA |
| 5 | Alix | CTGGAAGGATGCTTTCGATAAAGG | AGGCTGCACAATTGAACAACAC |
| 6 | MITF | AAGGGCTTGCAGAACACCTTA | GCTGGTTTGGACATGGCAAG |
| 7 | Rab27a | AGAGGAGGAAGCCATAGCAC | CATGACCATTTGATCGCACCAC |
| 8 | Rab27b | GGAACTGGCTGACAAATATGG | CAGTATCAGGGATTTGTGTCTT |
| 9 | GAPDH | CAAGGTCATCCATGACAACTTTG | GTCCACCACCCTGTTGCTGTAG |

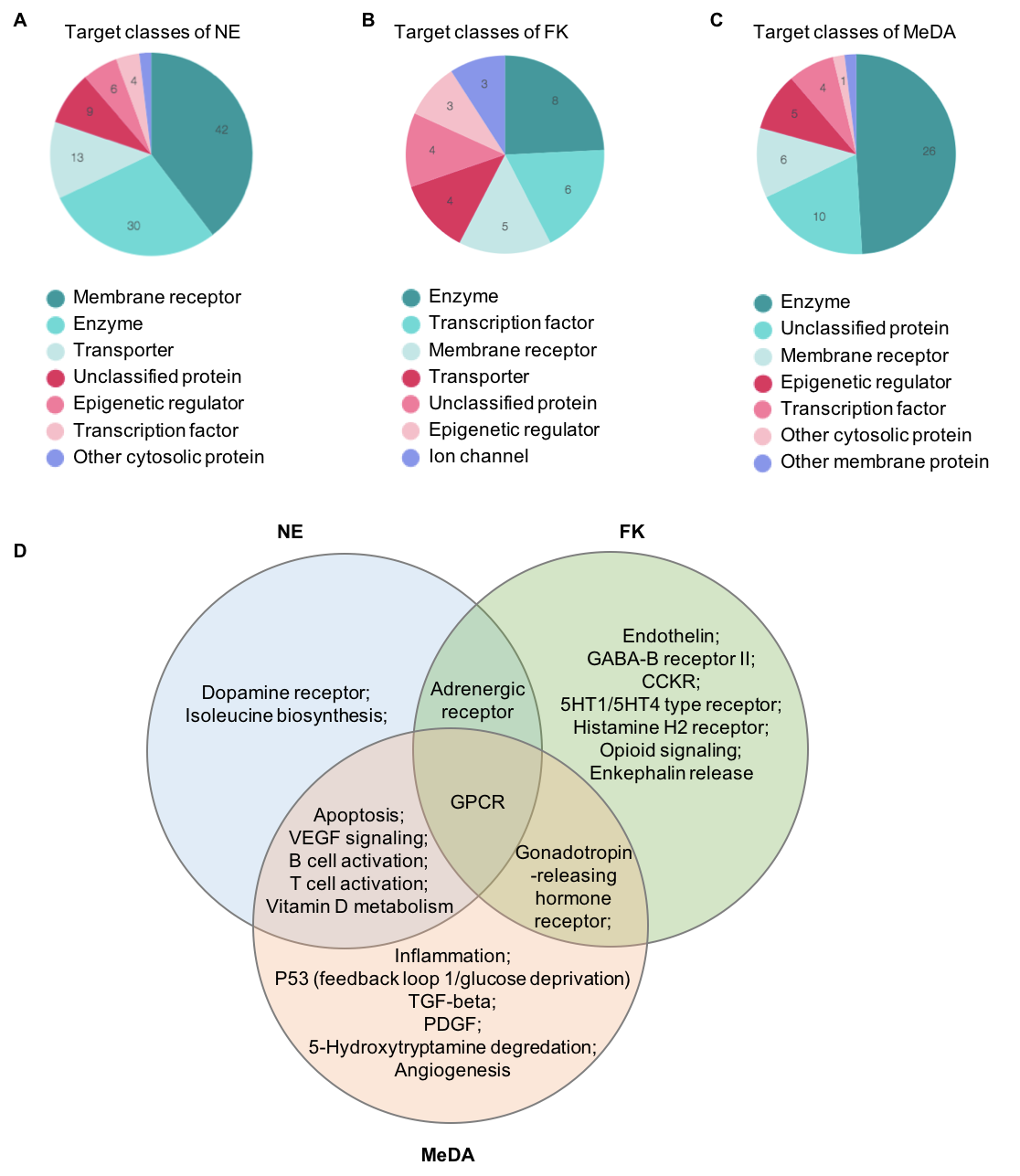

**Appendix Fig. 1: Target prediction of the compounds.** CheMBL database and SwissTargetPrediction were used to predict the potential targets of the tested compounds, (**A**) 42 targets of NE are membrane receptors and NE mainly targets different kinds of adrenergic receptors**.** (**B**) FK targets adenylate cyclase, glucagon-like peptide receptor, and glucose transporter, which are related to energy (ATP and glucose) production in cells. (**C**) MeDA targets dopamine receptors and inflammation related proteins.

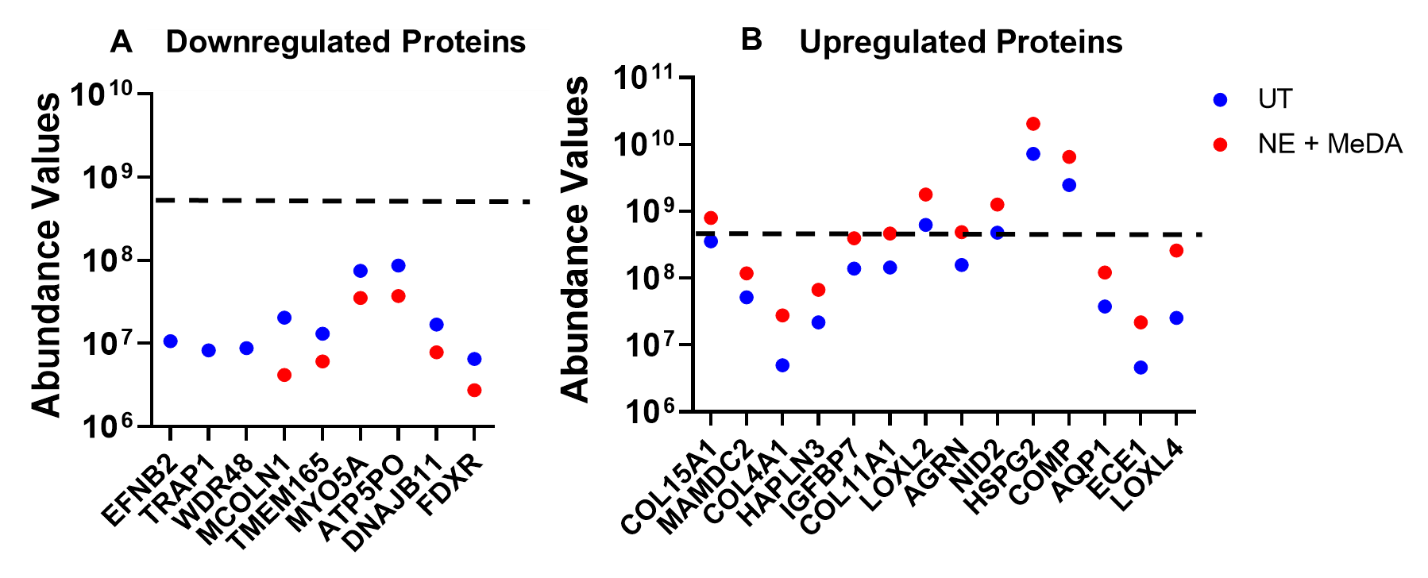

**Appendix Fig. 2: Assessment of protein abundance change with respect to average abundance value.** Proteins that were found to be up (A) or down (B) regulated between exosomes derived from non-treated MSCs (NT) and MSCs treated with NE and MeDA were plotted with respect to the average protein abundance (dotted line). Downregulated proteins were found to be relatively low abundance with respect to the average. Proteins that were upregulated over the average abundance (COL15A1, COL11A1, LOXL2, AGRN, NID2, HSPG2, COMP) were considered for pathway analysis.
